# Cultivation of *Melilotus officinalis* as a source of bioactive compounds in association with soil recovery practices

**DOI:** 10.1101/2023.04.24.538143

**Authors:** Isabel Nogues, Laura Passatore, María Ángeles Bustamante, Emanuele Pallozzi, João Luz, Francisco Traquete, António E.N. Ferreira, Marta Sousa Silva, Carlos Cordeiro

**Author notes:** Corresponding author: Marta Sousa Silva. **Author Contributions** I.N. and L.P. conceived the study, performed the experimental design and were involved in all analytical experiments. I.N., L.P., M.A.B. and E.P. performed plant growth assays, analytical determinations and gas-exchange measurements. I.N., L.P., M.S.S. and C.C. performed the untargeted metabolomics analysis. J.L., F.T. and A.E.N.F. developed the in-house software for metabolomics data analysis. J.L., M.S.S. and C.C. performed metabolomics data analysis and interpretation. I.N., L.P., M.S.S. and C.C. wrote the manuscript. All authors reviewed and approved the manuscript.

## Abstract

*Melilotus officinalis* is a Leguminosae with relevant applications in medicine and soil recovery. This study reports the application of *Melilotus officinalis* plants in soil recovery and as a source of bioactive compounds. Plants were cultivated in semiarid soil under four different fertilizer treatments, urban waste compost at 10 t/ha and 20 t/ha, inorganic fertilizer and a control (no fertilizer). Agronomic properties of soil (pH, EC, soil respiration, C content, macro- and micro-elements) were analyzed before and after treatment. Also, germination, biomass, element contents, and physiological response were evaluated. Results showed a significant enhancement of the soil microbial activity in planted soils amended with compost, though there were no other clear effects on the soil physicochemical and chemical characteristics during the short experimental period. An improvement in *M. officinalis* germination and growth was observed in soils with compost amendment. Metabolite composition of plants was analyzed through Fourier-transform ion cyclotron resonance mass spectrometry (FT-ICR MS). Principal Component and Agglomerative Hierarchical Clustering models suggest that there is a clear separation of the metabolome of four groups of plants grown under different soil treatments. The five most important discriminative metabolites (annotated) were oleamide, palmitic acid, stearic acid, 3-hydroxy-cis-5-octenoylcarnitine, and 6-hydroxynon-7-enoylcarnitine. This study provides information on how the metabolome of *Melilotus* might be altered by fertilizer application in poor soil regions. These metabolome changes might have repercussions for the application of this plant in medicine and pharmacology. The results support the profitability of *Melilotus officinalis* cultivation for bioactive compounds production in association with soil recovery practices.

## Introduction

*Melilotus officinalis* is a common biannual Leguminosae known as yellow sweet clover. *M. officinalis* is a pioneer herb, native to Eurasia and naturalized in Europe throughout Asia and America. This plant has important applications in two relevant fields: medicine and soil recovery. Concerning medicinal applications, biological activity studies have demonstrated the antioxidant, anti-inflammatory, and antiproliferative effects of *M. officinalis* (Trouillas et al., 2003; Miliauskas et al., 2004). The European Medicines Agency reported using *M. officinalis* orally to relieve discomfort and heaviness of legs related to minor venous circulatory disturbances and topically for minor inflammations of the skin (EMA, 2017). Studies showed that this plant prevented skin ageing, promoted tissue regeneration, and reduced fat deposition (Pastorino et al., 2017). Moreover, ANGIPARS™ (semelil), an herbal drug containing *Melilotus officinalis* extract and produced in oral, topical, and intravenous forms, has been shown to have a positive effect in the treatment of diabetic foot ulcers (Chorepsima et al., 2013). Semelil is mainly composed of coumarin derivatives, flavonoids, and selenium (Aslroosta et al., 2021). Studies about the phytochemical profile of *M. officinalis* reported that plant tissues contain flavonoids such as quercetin derivatives and various phenolic compounds, coumarins, steroids and saponins (Anwer et al., 2008; Tang, 2012; Yang et al., 2014).

Nowadays, soil health, functionality and productivity are severely hampered by continuous crop production, clearing of vegetation, urbanization, adverse climatic conditions (erratic rainfall, floods, periodic droughts) and forest fires (Gomez-Sagasti et al. 2018; Olsson et al., 2019). All these factors contribute to the loss of soil organic matter (SOM), which is strongly linked to soil degradation and desertification, a serious problem not only in terms of agricultural productivity but also for the reclaiming of landscapes for recreation and nature conservation. Within this context, revegetation is an effective tool for improving the quality of soil and rehabilitating degraded environments (Zhang et al., 2011; Gao et al., 2018) as it enhances the storage of SOM in soils by reducing soil erosion and increasing inputs of organic materials. The use of profitable crops for soil recovery strategies leads to a net gain in soil health and functionality, generating at the same time revenues for landowners during the soil restoration treatment. In this sense, the ability of *M. officinalis* to develop root nodules and fix nitrogen in symbiosis with rhizospheric bacteria, even in saline and arid environments, makes this plant useful for soil recovery purposes and for the reduction of large inorganic N fertilizer needs (Houérou, 2001; Song et al., 2022).

Moreover, positive effects on soil quality may be enhanced by the application to the soil of organic amendments (Barra-Caracciolo et al., 2015; Tully and McAskill, 2020). Indeed, the addition of composts is essential for the recovery of the carbon (C) and nitrogen (N) stocks, especially in arid and semiarid soils, since it maintains soil organic matter levels, supplies nutrients and enhances microbial proliferation and activity (Tejada et al., 2006). In this context, the use of municipal waste-derived compost may contribute to soil fertility recovery, allowing at the same time environment-friendly management of the organic fraction of municipal solid waste by reducing the amount of organic waste directed to incineration or landfill disposal.

The present study aimed to investigate the cultivation of *M. officinalis* for different purposes: as a cover plant for soil recovery and as a profitable cultivation with bioactive compounds for the pharmacological industry. Physiological analyses were carried out on plants to evaluate the plant response to different nutritive conditions and its adaptability to grow in poor soils. Soil characterization was performed before and after the treatment to assess the efficiency of each treatment for soil recovery. Fourier Transform Ion Cyclotron Resonance Mass Spectrometry (FT-ICR-MS) was used for the metabolic profiling of *M. officinalis* and to understand the effect of different fertilizing treatments on its metabolome and to validate the profitability of *Melilotus officinalis* cultivation for bioactive compounds production in association with soil recovery practices.

## Results

### Changes on the soil physico-chemical, chemical and biological characteristics

The incorporation of the compost and the inorganic fertilizer in the soil obtained from a semiarid abandoned agricultural area, did not modify soil pH, most likely due to the buffering effect of the alkaline and calcareous soil (**Table 1**). Soil electrical conductivity (EC) values did not present significant differences among treatments six months after the experimental setup (**Table 1**). However, the presence of plants for three months caused some modifications. In Control soils, the amount of salt in the soil decreased three months after sowing, whereas in inorganic fertilized soil, the opposite effect was observed. Indeed, at the end of the experiment inorganic fertilized soil presented the highest EC value, 2-, 1.8- and 1.6-fold higher than the control, C-Low and C-High, respectively.

**Table 1:** Soil pH, EC and respiration in Control, Inorganic, C-Low and C-High treatments at the beginning (prior to plant sowing) and at the end (three months after sowing). Different letters mean significant differences among treatments at the same experimental time (n=5). Asterisks mean significant differences between values at the beginning and at the end for each individual treatment.

| Parameter |  | Control | Inorganic | C-Low | C-High |
| --- | --- | --- | --- | --- | --- |
| pH | Beginning | 8.07a ± 0.04 | 8.13a ± 0.01 | 8.04a ± 0.07 | 7.99a ± 0.01 |
|  | End | 8.13a ± 0.08 | 8.02a ± 0.06 | 8.23a ± 0.08 | 7.94a ± 0.03 |
| EC (dS/m) | Beginning | 0.81a ± 0.05 | 0.72a ± 0.05 | 0.88a ± 0.10 | 0.77a ± 0.02 |
|  | End | 0.53b ± 0.02 | 1.04a ± 0.07 | 0.81b ± 0.10 | 0.63b ± 0.04 |
|  |  | * | * |  |  |
| Soil respiration<br>(mg CO<sub>2</sub>-C/kg<br>d.w.·day) | Beginning | 28.27d ± 5.24 | 93.21a ± 7.81 | 63.10b ± 5.11 | 51.35c ± 3.26 |
|  | End | 37.68c ± 5.87 | 64.70a ± 7.57 | 77.76a ± 5.70 | 72.73b ± 5.23 |
|  |  |  | * |  | * |

However, unlike pH and EC performances, soil respiration values were different between treatments already at the first sampling point, showing the soil with the mineral fertilization (Inorganic) a value of this parameter 3.3-, 1.5- and 1.8-fold higher than Control, C-Low and C-High, respectively (**Table 1**). This fact could be due to the presence of easily available nutrients in the treatment with inorganic fertilization, which could have initially enhanced the activity of the microorganisms, which quickly reduced after three months from sowing. The treatments with compost, C-Low and C-High with the presence of the plant also presented high soil respiration values. However, Control soil without fertilization, either before or three months after yellow clover plants sowing, presented the lowest respiration values. Soil respiration value of Control soils was 1.7-, 2.1-, 1.9-fold lower than Inorganic, C-Low and C-High soils, respectively.

The total Nitrogen (N_T_) and total Carbon (C_T_) rates in soil were measured in the different experimental conditions at the beginning and the end of the experimental set-up (**Figure 1, A and B**). At the beginning, soil N_T_ resulted to be 25%, and 51% higher in the soils treated with the organic amendment at the two different doses compared to the control soil. However, N_T_ in the inorganic-fertilized soil was only slightly higher than in the non-fertilized one (8% higher). At the end of the experiment, the difference between Control and the C-High soil increased further: N_T_ resulted to be 20% and 70% higher in C-Low and C-High soils than in the control soil, respectively. Furthermore, N_T_ in the inorganic-fertilized soil was only slightly higher than in the control one, as measured at the beginning of the experiment (10%).

**Figure 1:**
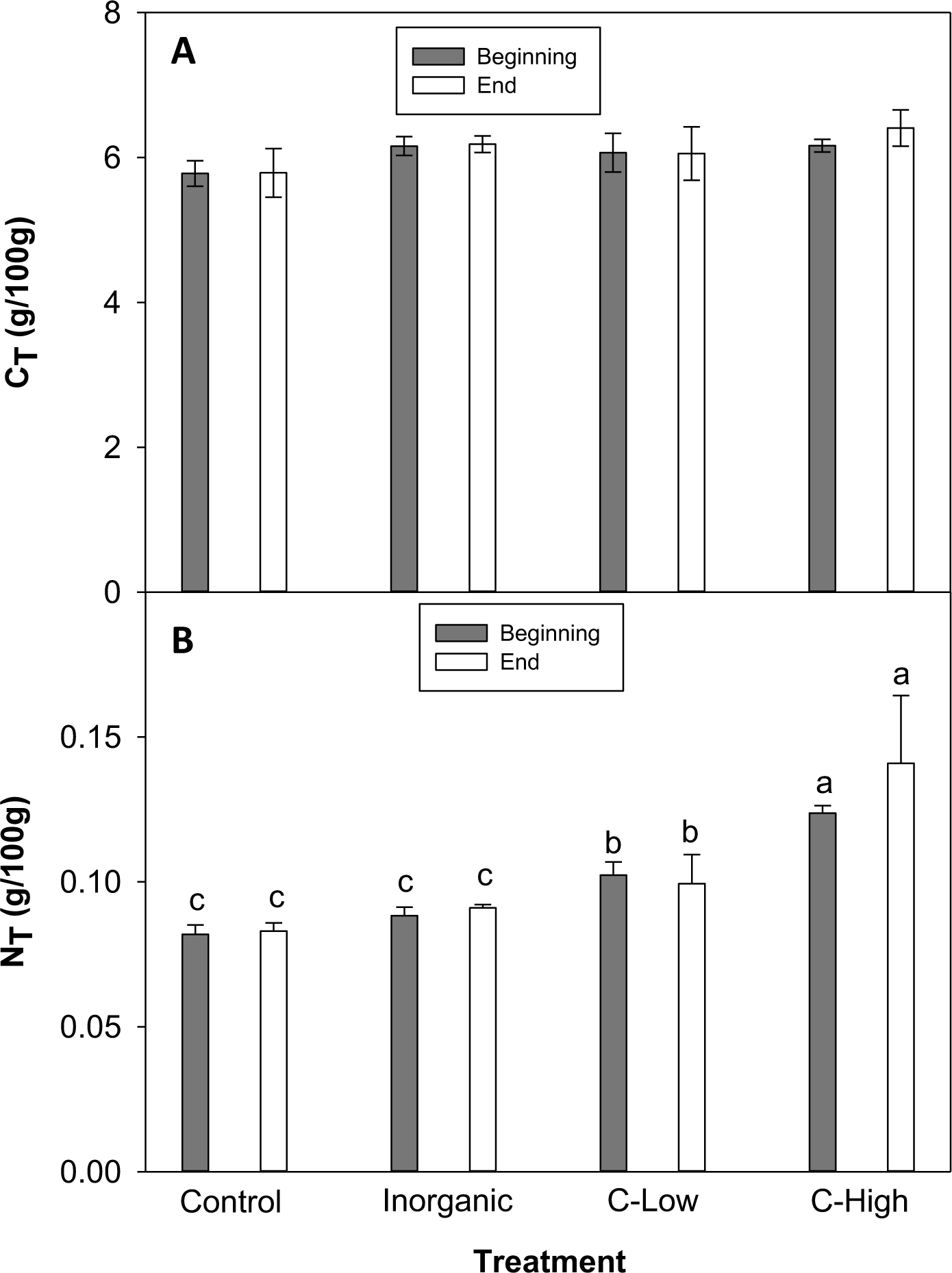
**A)** Soil C_T_ and **B)** Soil N_T_ contents (g/100mg) in Control, Inorganic, C-Low and C-High treatments at the beginning (prior to plant sowing) and at the end (three months after sowing). Different letters mean significant differences among treatments at the same experimental time (n=5).

Concerning macroelements in soils, we did not find any significant difference in P, Ca, Mg values among treatments before *M. officinalis* sowing (**Table 2**). At the end of the experimental period, however, Mg values resulted to be higher in the control soil than in the inorganic and C-High soils. On the other hand, K and Na contents were higher in compost-amended soils than in inorganic fertilized soil at the beginning of the experiment. At the end, there were no significant differences among treatments (**Table 2**).

**Table 2:**
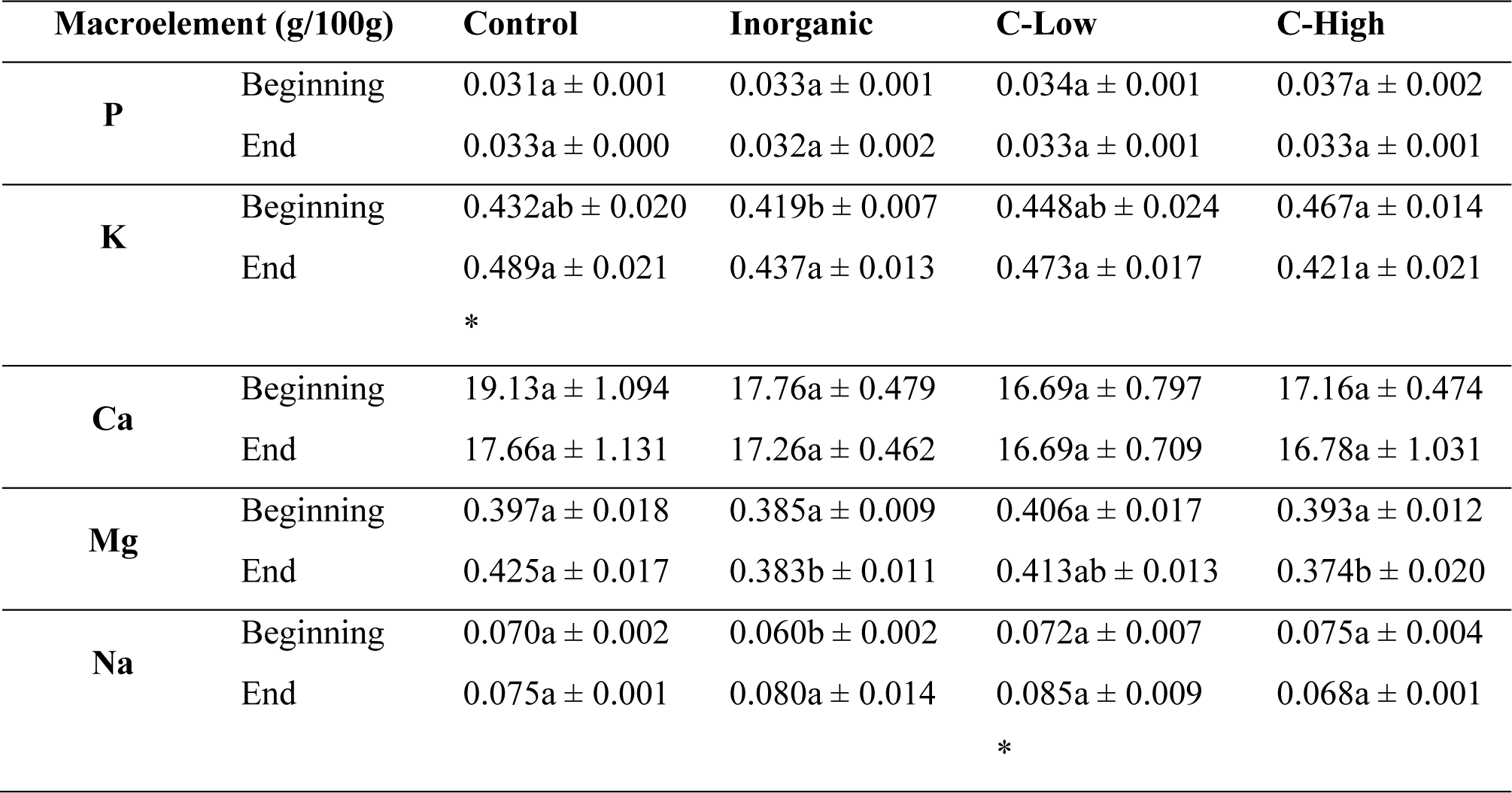
Soil P, K, Ca Mg and Na contents (g/100mg) in Control, Inorganic, C-Low and C-High treatments at the beginning (prior to plant sowing) and at the end (three months after sowing). Different letters mean significant differences among treatments at the same experimental time (n=5). Asterisks mean significant differences between values at the beginning and at the end for each individual treatment.

| Macroelement (g/100g) |  | Control | Inorganic | C-Low | C-High |
| --- | --- | --- | --- | --- | --- |
| P | Beginning | 0.031a ± 0.001 | 0.033a ± 0.001 | 0.034a ± 0.001 | 0.037a ± 0.002 |
|  | End | 0.033a ± 0.000 | 0.032a ± 0.002 | 0.033a ± 0.001 | 0.033a ± 0.001 |
| K | Beginning | 0.432ab ± 0.020 | 0.419b ± 0.007 | 0.448ab ± 0.024 | 0.467a ± 0.014 |
|  | End | 0.489a ± 0.021 | 0.437a ± 0.013 | 0.473a ± 0.017 | 0.421a ± 0.021 |
|  |  | * |  |  |  |
| Ca | Beginning | 19.13a ± 1.094 | 17.76a ± 0.479 | 16.69a ± 0.797 | 17.16a ± 0.474 |
|  | End | 17.66a ± 1.131 | 17.26a ± 0.462 | 16.69a ± 0.709 | 16.78a ± 1.031 |
| Mg | Beginning | 0.397a ± 0.018 | 0.385a ± 0.009 | 0.406a ± 0.017 | 0.393a ± 0.012 |
|  | End | 0.425a ± 0.017 | 0.383b ± 0.011 | 0.413ab ± 0.013 | 0.374b ± 0.020 |
| Na | Beginning | 0.070a ± 0.002 | 0.060b ± 0.002 | 0.072a ± 0.007 | 0.075a ± 0.004 |
|  | End | 0.075a ± 0.001 | 0.080a ± 0.014 | 0.085a ± 0.009 | 0.068a ± 0.001 |
|  |  |  |  | * |  |

Concerning microelements, the situation was similar (**Table 3**). No significant differences among treatments were found for Fe, Cd, Cu, Mn, Ni and Pb contents at both sampling times and for Mo and Zn only at the final sampling. On the other hand, at time 0, Mo resulted to be higher in Control and Inorganic soils than in compost-amended ones, contrarily to Zn, which was higher in C-Low and C-High soils (**Table 3**).

**Table 3:** Soil Fe, Cd, Cu, Mo, Mn, Ni, Pb and Zn contents (mg/kg) in Control, Inorganic, C-Low and C-High treatments at the beginning (prior to plant sowing) and at the end (three months after sowing). Different letters mean significant differences among treatments at the same experimental time (n=5). Asterisks mean significant differences between values at the beginning and at the end for each individual treatment.

| Microelement (mg/kg) |  | Control | Inorganic | C-Low | C-High |
| --- | --- | --- | --- | --- | --- |
| **Fe** | Beginning | 17.46a ± 1.37 | 15.14a ± 0.41 | 15.67a ± 1.14 | 14.98a ± 0.31 |
|  | End | 17.09a ± 1.37 | 14.47a ± 0.48 | 15.62a ± 1.06 | 14.54a ± 0.62 |
| **Cd** | Beginning | 0.451a ± 0.028 | 0.610a ± 0.128 | 0.472a ± 0.021 | 0.457a ± 0.010 |
|  | End | 0.496a ± 0.035 | 0.439a ± 0.019 | 0.479a ± 0.019 | 0.431a ± 0.020 |
| **Cu** | Beginning | 9.02a ± 0.548 | 8.66a ± 0.200 | 9.35a ± 0.394 | 9.77a ± 0.36 |
|  | End | 9.87a ± 0.741 | 8.52a ± 0.233 | 9.39a ± 0.379 | 8.98a ± 0.25 |
| **Mo** | Beginning | 0.410a ± 0.01 | 0.556ab ± 0.256 | 0.378ab ± 0.033 | 0.314b ± 0.020 |
|  | End | 0.309a ± 0.016 | 0.332a ± 0.042 | 0.312a ± 0.015 | 0.258a ± 0.005 |
|  |  | * |  |  |  |
| **Mn** | Beginning | 305.75a ± 23.21 | 295.62a ± 2.99 | 309.32a ± 14.50 | 286.29a ± 6.32 |
|  | End | 321.58a ± 11.92 | 284.59a ± 15.31 | 306.17a ± 8.78 | 278.02a ± 13.05 |
| **Ni** | Beginning | 18.69a ± 1.759 | 17.57a ± 0.111 | 19.09a ± 1.151 | 17.23a ± 0.41 |
|  | End | 20.35a ± 1.559 | 17.08a ± 0.888 | 18.82a ± 0.798 | 16.93a ± 0.37 |
| **Pb** | Beginning | 13.71a ± 0.451 | 13.30a ± 0.102 | 14.48a ± 0.999 | 13.88a ± 0.431 |
|  | End | 15.56a ± 1.07 | 13.52a ± 0.605 | 14.35a ± 0.715 | 13.18a ± 0.353 |
| **Zn** | Beginning | 34.62ab ± 1.980 | 33.53b ± 0.409 | 36.19ab ± 1.904 | 36.24a ± 0.843 |
|  | End | 37.46a ± 2.114 | 32.51a ± 1.201 | 35.87a ± 1.274 | 33.99a ± 1.100 |

### Effects on plant characteristics: germination, yield, element contents and physiological response

Maximum germination was obtained in high compost amended soils (71%) and in inorganic fertilized soils (68%). The percentage of germinated seeds was similar in control and in low-dose composted soils, 59% and 57% respectively (**Figure 2A**). Concerning the germination rate (**Figure 2B**) it resulted also to be higher in high-compost amended soils (C-High) and in inorganic fertilized soils (Inorg). Noticeably, the lowest germination rate was found in low-compost amended soils (C-Low).

**Figure 2:**
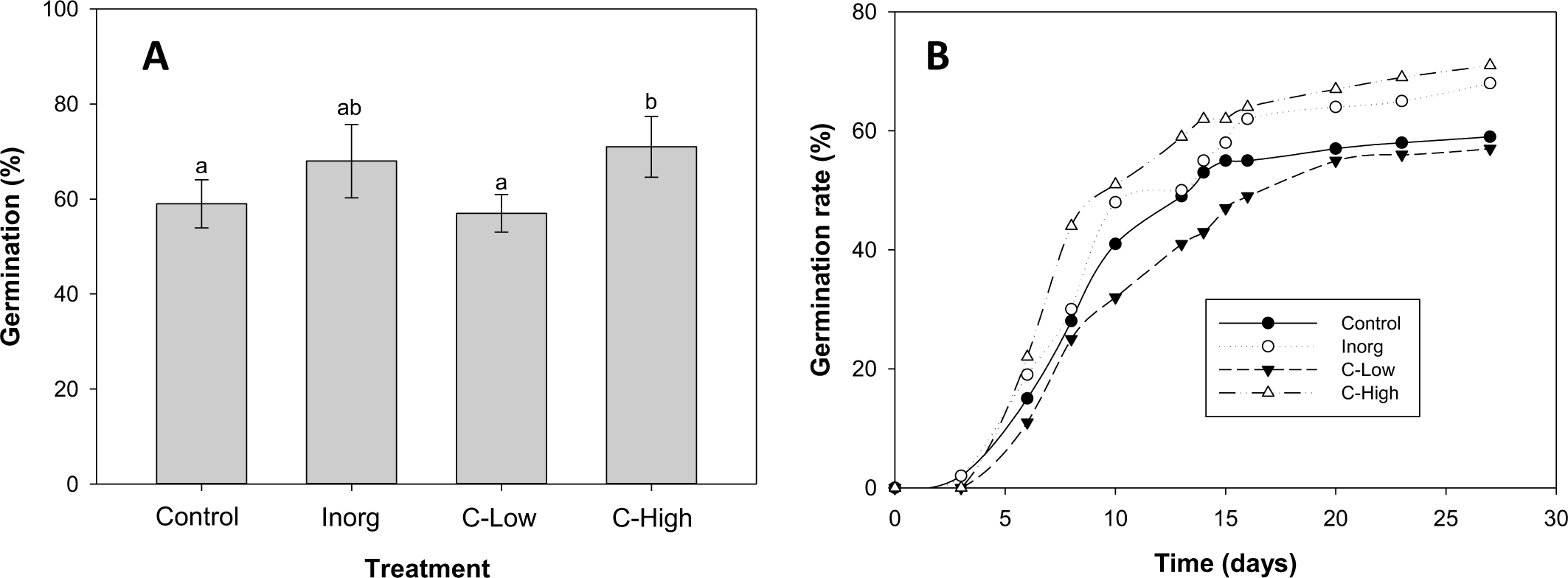
**A)** Germination percentage (left axis) and Maguires germination rate index (right axis) **B)** Emergency rate of *Melilotus officinalis* seeds sowed on Control, Inorganic, C-Low and C-High soils. Different letters mean significant differences among treatments (n=10).

The average values of *M. officinalis* root plant dry biomass were significantly lower (p<0.001) in the Control than in treatments with compost-amended soils, 1.6- and 2.5-fold lower, respectively (p<0.001) (Fig. 3A). Moreover, root biomass was higher in C-High plants than in C-Low ones.

**Figure 3:**
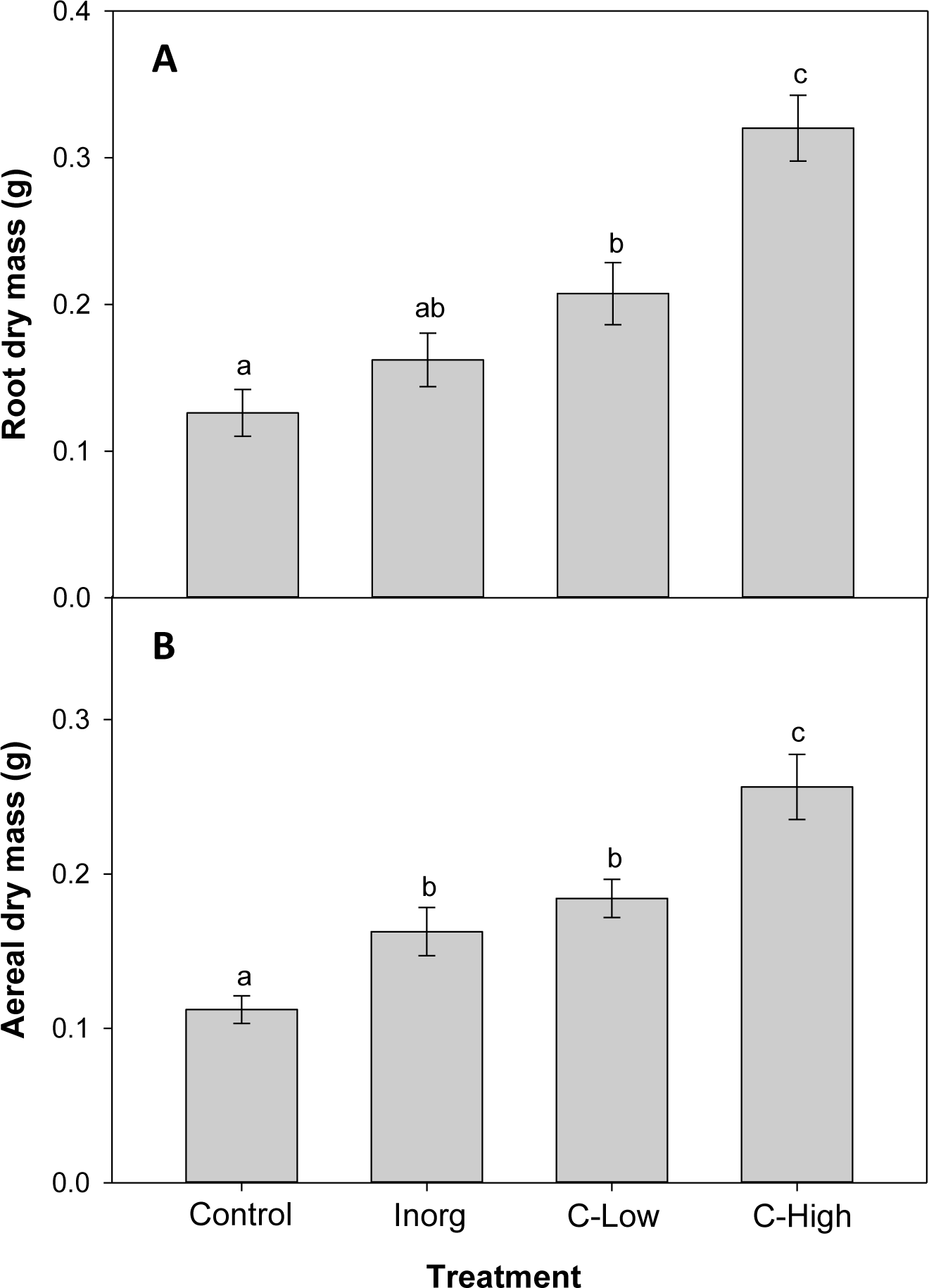
**A)** Root dry biomass and **B)** Aerial dry biomass of *Melilotus officinalis* plants grown on Control, Inorganic, C-Low and C-High soils. Different letters mean significant differences among treatments (n=10).

Concerning aerial dry biomass, values were 1.4-, 1.6- and 2.3-fold lower (p<0.001) in Control than in inorganic, C-Low and C-High treatments, respectively (**Figure 3B**). Moreover, amendment with high dose compost increased the plant aerial biomass 1.6-fold with respect to inorganic fertilizer and 1.4-fold in comparison with low-dose compost.

We also determined root lengths, that resulted to be very different among soil treatments: Control 19.98 ± 2.96; Inorganic 14.54 ± 6.97; C-Low 20.22 ± 8.13; C-High 30.52 ± 20.10, also reflecting differences in root morphology (**Figure 4**). In all treatments, roots formed nodules for N_2_ fixation, typical of the *M. officinalis* species.

**Figure 4:**
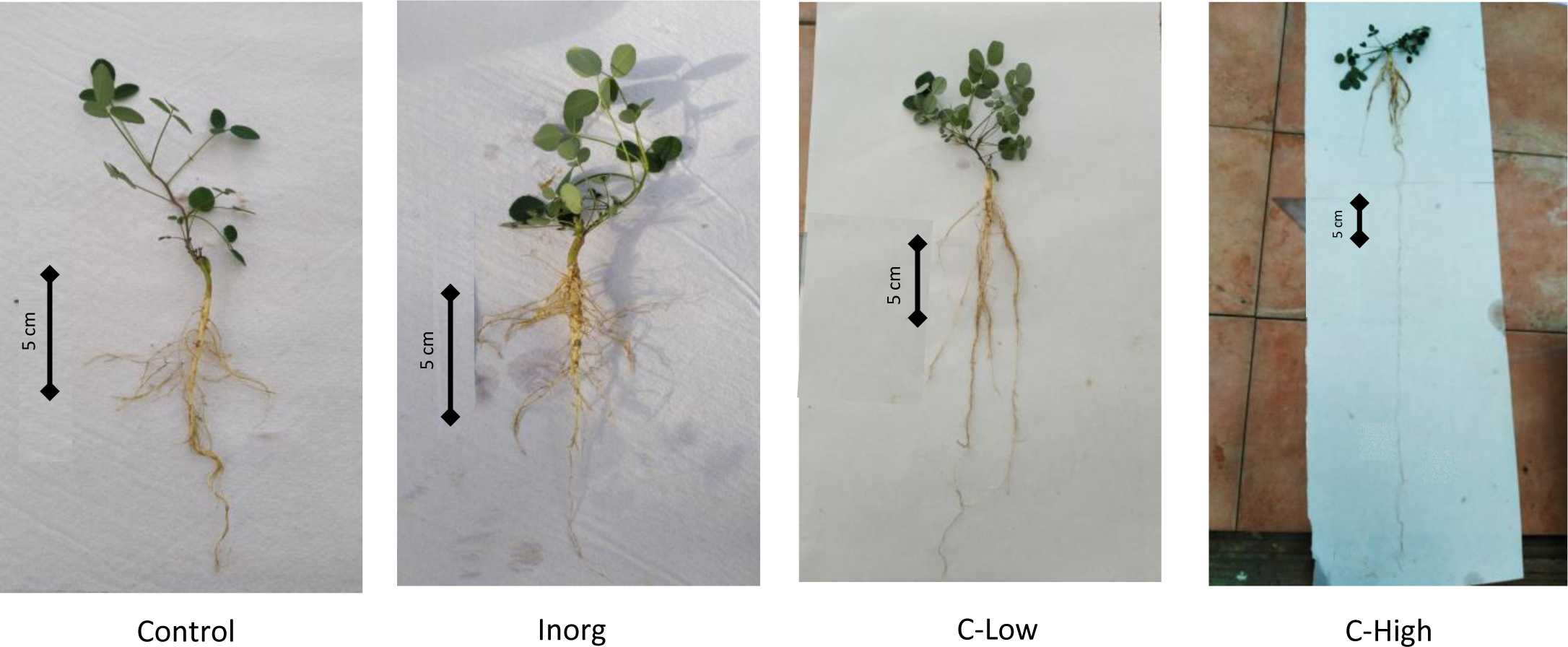
Pictures of the plants sampled at the end of the treatment. One plant per treatment has been chosen as representative.

No differences among treatments were found for gas-exchange response at the leaf level (**Table 4**). CO_2_ assimilation was in the range of 26 – 28 µmol m^-2^ s^-1^ according to previously reported values for *M. officinalis* plants (Peterson et al., 2004). Stomatal conductance ranged between 0.484 and 0.592 mmol m^-2^ s^-1^ among the different treatments.

**Table 4:** Leaf gas exchange parameters of *Melilotus officinalis* plants grown on Control, Inorganic, C-Low and C-High soils (n=5).

| Treatment | Photosynthesis ($\mu\text{mol m}^{-2}\text{s}^{-1}$) | Stomatal conductance ($\text{mmol m}^{-2}\text{s}^{-1}$) |
| --- | --- | --- |
| Control | 26.39 ± 1.95 | 0.592 ± 0.085 |
| Inorganic | 27.81 ± 1.45 | 0.484 ± 0.073 |
| C-Low | 27.94 ± 2.08 | 0.532 ± 0.099 |
| C-High | 27.79 ± 1.89 | 0.493 ± 0.051 |

Total Nitrogen and total Carbon values were measured in the leaves of *Melilotus officinalis* plants (**Figure 5**). N_T_ values resulted to be c.a 10% higher in leaves of plants grown in soils amended with compost, independently of the dose used, than in any other experimental condition. Leaf C_T_, however, resulted to be similar in all treatments (**Figure 5**).

**Figure 5:**
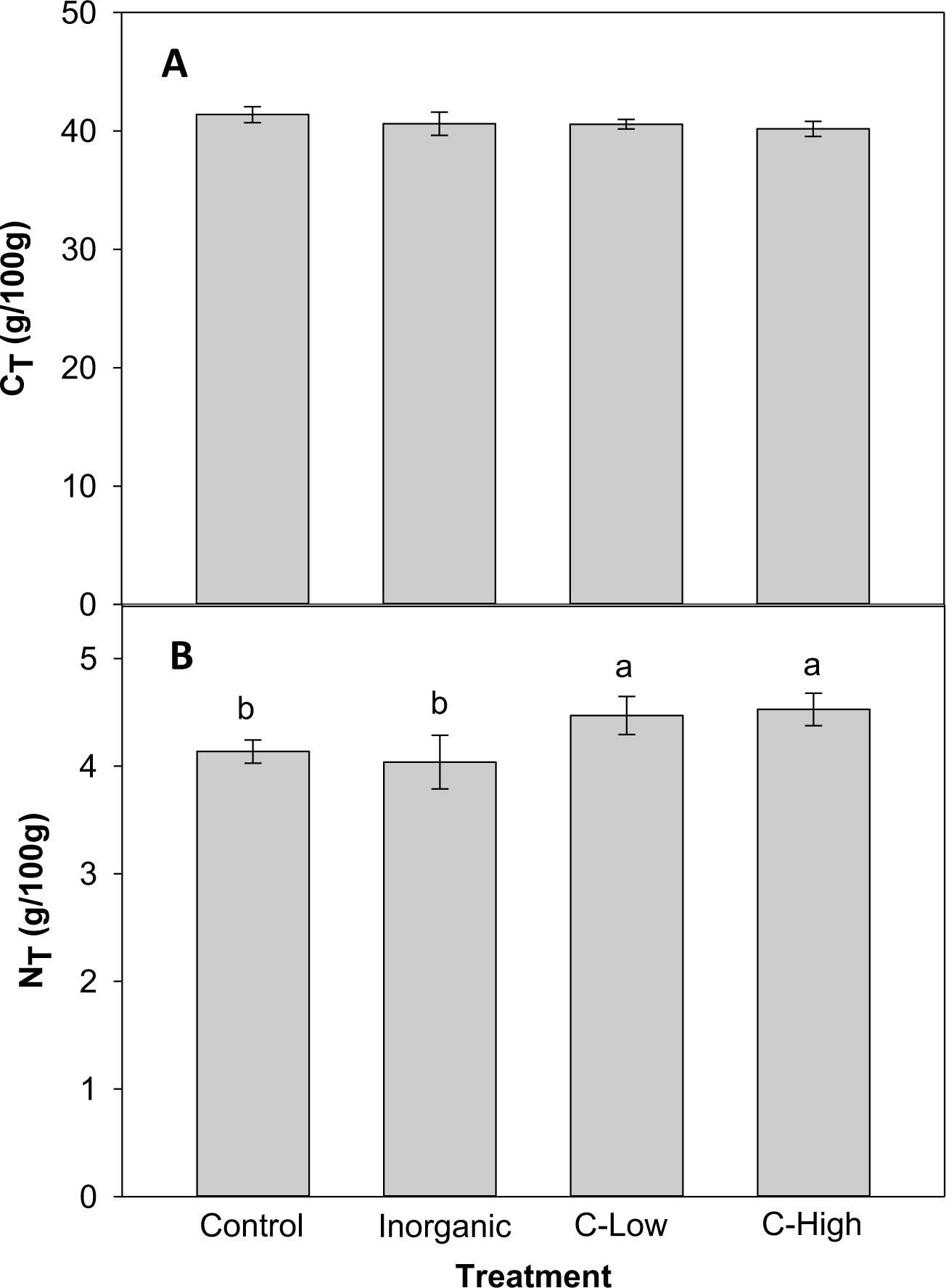
**A)** Leaf C_T_ and **B)** Leaf N_T_ contents (g/100mg) in Control, Inorganic, C-Low and C-High treatments. Different letters mean significant differences among treatments (n=5).

The contents of macro- and microelements were determined in 3 months old *M. officinalis* leaves (**Tables 5 and 6**). It is interesting to know that, whereas leaf P content was higher in plants grown on compost-amended soils than in plants grown on inorganic fertilized or control soils, leaf K, Ca, Mg and Na behaved in the other way round. In general, also Cd, Mn, Ni and Zn tended to be lower in C-Low and C-High plants than in inorganic and control plants (**Table 6**). On the other hand, leaf Mo, Cu, and Pb contents were similar in all treatments, whereas Fe resulted to be lower in leaves grown in Inorganic soil than in Control, C-Low and C-High leaves.

**Table 5:** Leaf P, K, Ca Mg and Na contents (g/100mg) in Control, Inorganic, C-Low and C-High treatments. Different letters mean significant differences among treatments (n=5).

| Macroelement<br>(g/100 g) | Control | Inorganic | C-Low | C-High |
| --- | --- | --- | --- | --- |
| P | 0.209b ± 0.016 | 0.229b ± 0.024 | 0.262a ± 0.020 | 0.290a ± 0.014 |
| K | 1.002a ± 0.143 | 0.800a ± 0.178 | 0.323b ± 0.053 | 0.263b ± 0.046 |
| Ca | 4.770a ± 0.731 | 5.335a ± 1.269 | 2.729b ± 0.199 | 2.687b ± 0.297 |
| Mg | 0.411a ± 0.105 | 0.347a ± 0.111 | 0.118ab ± 0.032 | 0.086b ± 0.019 |
| Na | 0.121a ± 0.024 | 0.078ab ± 0.028 | 0.059ab ± 0.015 | 0.029b ± 0.005 |

**Table 6:** Leaf Fe, Cd, Cu, Mo, Mn, Ni, Pb and Zn contents (mg/Kg) in Control, Inorganic, C-Low and C-High treatments. Different letters mean significant differences among treatments (n=5).

| Microelement<br>(mg/kg) | Control | Inorganic | C-Low | C-High |
| --- | --- | --- | --- | --- |
| Fe | 237.4a ± 14.15 | 176.86b ± 17.30 | 260.02a ± 11.98 | 258.59a ± 45.19 |
| Cd | 0.738a ± 0.426 | 0.2378a ± 0.1083 | 0.0615b ± 0.0109 | 0.0779b ± 0.0209 |
| Cu | 22.40a ± 5.440 | 12.480a ± 2.667 | 11.157a ± 0.668 | 10.698a ± 0.307 |
| Mo | 5.366a ± 2.154 | 3.116a ± 1.351 | 4.786a ± 1.694 | 3.919a ± 0.900 |
| Mn | 164.1a ± 8.050 | 148.91ab ± 13.53 | 123.48b ± 9.19 | 117.47b ± 8.61 |
| Ni | 2.824a ± 1.002 | 1.545ab ± 0.397 | 0.899b ± 0.083 | 1.245ab ± 0.581 |
| Pb | 4.086a ± 1.183 | 3.895a ± 0.773 | 4.011a ± 0.906 | 2.359a ± 0.589 |
| Zn | 113.0a ± 34.95 | 35.42b ± 4.57 | 35.46b ± 2.91 | 31.36b ± 1.69 |

### FT-ICR MS metabolic analysis in leaves

An untargeted metabolomics analysis using FT-ICR-MS, with electrospray ionization in positive (ESI^+^) mode was performed. A total of 4490 putative metabolites were detected and, within each of the four soil classes Control, Inorg, C-High and C-Low, the number of compounds was similar (**Figure 6A**). To gain insight into metabolite diversity, chemical formulas were predicted for 1471 metabolites by using SmartFormula. Moreover, 1019 metabolites were identified by name with HMDB, PlantCyc, and LOTUS. A total of 406 annotated (by name) metabolites were common to all samples, as shown in the intersection plot (**Figure 6A**). Among them, we found bioactive compounds such as coumarin, melilotoside, o-coumaric acid or p-coumaric acid, and carvone (already described in *Melilotus* plants according to the European Medicines Agency Assessment report on *Melilotus officinalis* (L.) Lam., EMA, 2017), ethyl palmitate, and dodecanoic acid/lauric acid. All identified and annotated compounds can be found in **Supplementary Table S1**. The control group had the highest number of exclusive metabolites, whereas the inorganic amendment group has the lowest number of exclusive metabolites. Under the ChemOnt classification, most of the compounds belong to the lipids and lipid-like molecules superclass, followed by organic acids and derivatives (**Figure 6B**). Within the different lipids classes, fatty acyls are most represented, followed by prenol lipids and steroids (**Figure 6B** **inset**). Examples include palmitic acid, α-linolenic acid, oleic acid, oleamide, linoleamide, 3-oxohexadecanoic acid, tridecanoic acid, among others (**Supplementary Table S1**).

**Figure 6:**
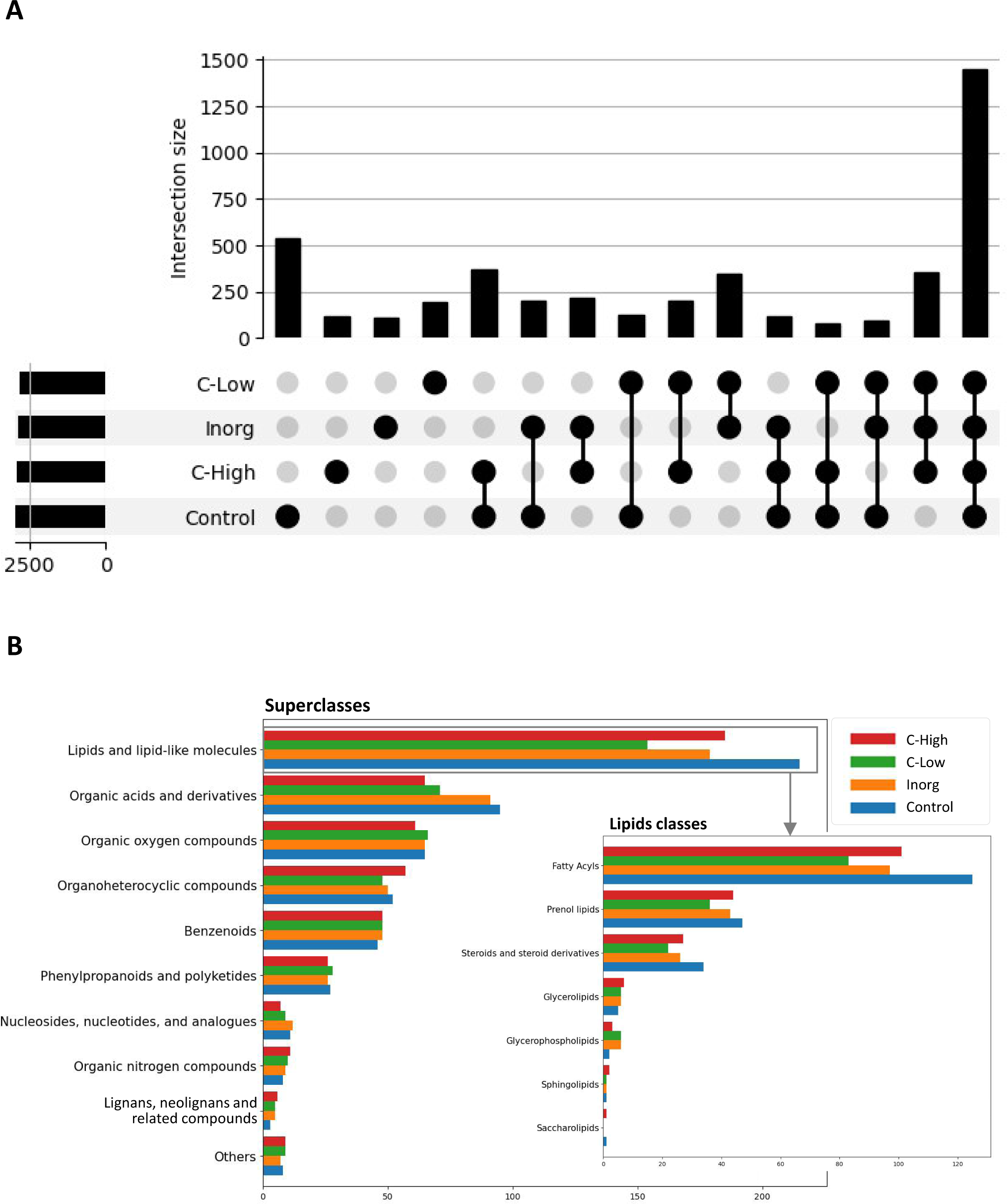
**A)** Intersection plot with common and exclusive metabolites among leaves in the four treatment groups; **B)** Compound classification into superclasses, with the insert showing the compound distribution within the different lipids’ classes.

Principal component analysis and hierarchical clustering were used to evaluate the analytical reproducibility and to ascertain the metabolic profile similarities between treatments (**Figure 7, A and B**). A clear separation between the control samples and the soil-treated samples was observed in the PCA scores plot, and, to a lesser extent, between the remaining groups along PC1 (**Figure 7A**). Hierarchical clustering showed the same trend, a clear separation of the control, with all replicates clustering closer together (**Figure 7B**). Indeed, the dendrograms in the hierarchical clustering were separated in two major clusters one containing control samples and the other with the rest of treatments, indicating that the metabolome profiling of *M. officinalis* leaves is sufficiently sensitive to distinguish unfertilized from fertilized grown plants (**Figure 7B**). The C-High and Inorg samples showed a higher similarity than the samples grown in C-Low. A supervised PLS-DA model was built for the purpose of classification and variable importance scoring (**Figure 8**). The significance of this model was assessed by a permutation test and a *p*-value below 0.05 was obtained. The VIP score for each variable was calculated to reveal the ones that most contribute to group discrimination in the PLS-DA model. The five most important discriminative metabolites (annotated) were oleamide, palmitic acid (hexadecanoic), stearic acid (octadecanoic acid), 3-hydroxy-cis-5-octenoylcarnitine, and 6-hydroxynon-7-enoylcarnitine. Their relative abundances in the different samples are shown in **Figure 9**. Oleamide levels were higher in the C-High-supplemented plants, as well as palmitoleamide and linoleamide, also discriminating compounds in the top 20 and top 40 discriminating compounds, respectively. Palmitic acid, stearic acid, 3-hydroxyhydroxy-cis-5-octenoylcarnitine and 6-hydroxynon-7-enoylcarnitine were more abundant in the control plants, with lower levels in C-Low.

**Figure 7:**
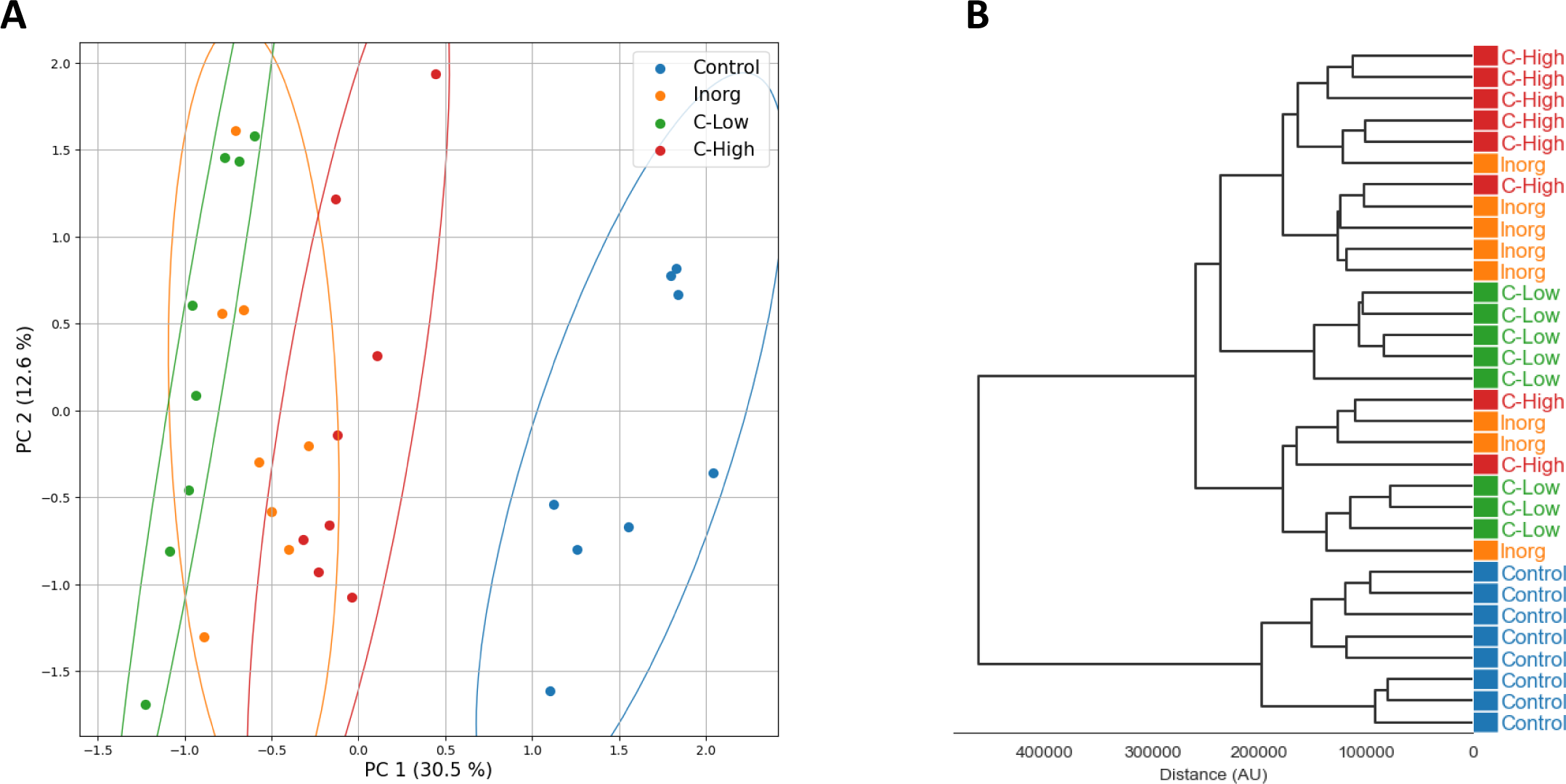
**A)** Principal component analysis (PCA) and **B)** hierarchical clustering analysis (HCA) of untargeted metabolomics obtained in positive electrospray ionization mode (ESI^+^).

**Figure 8:**
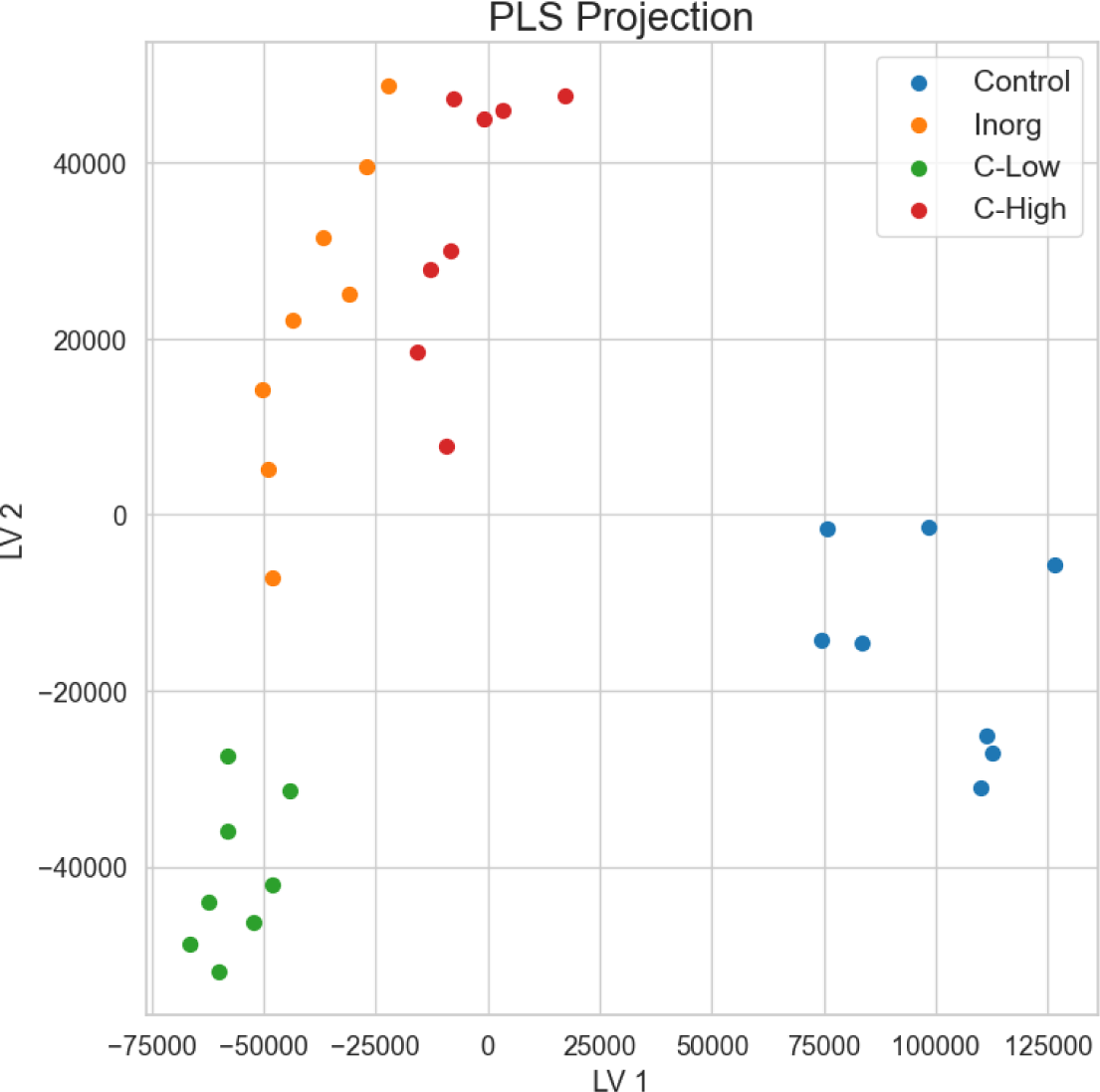
Partial least squares discriminant analysis (PLS-DA) for the separation of the four soil treatments.

**Figure 9:**
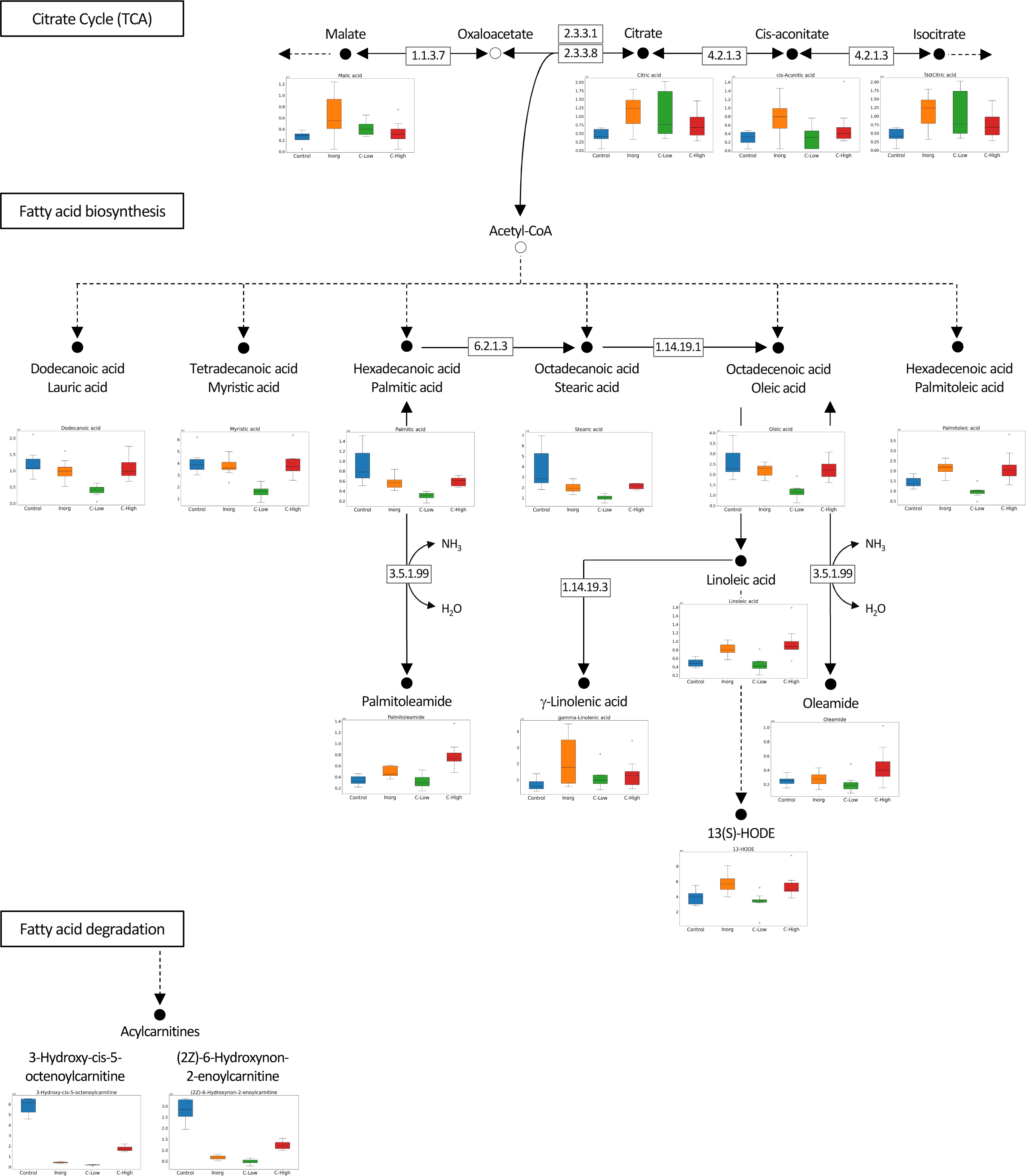
Metabolic pathway analysis evidencing reactions from the citrate cycle (TCA), fatty acid biosynthesis, and fatty acid degradation. Relative abundances of several discriminating compounds in *M. officinalis* plants grown on Control, Inorganic, C-Low and C-High soils, are represented as box plots.

## Discussion

### Responses of soil properties to soil amendment treatments

Soil properties, electrical conductivity (EC), pH and soil respiration responded differently to soil treatments and/or to the presence of plants. Whereas soil pH did not change with fertilization treatments, we observed changes in soil EC as a result of the combination between plant and type of fertilization. In control soils, plants reduced soil EC, but in inorganic-amended soils plants had the opposite effect. Root activities, such as transporting ions, releasing exudates and changing soil structure, may have a large influence on soil EC (Cao et al., 2010). In our case, *M. officinalis* root activities responded differently to different types of fertilization. Soil respiration is a measure of the carbon dioxide released from the soil by microbes decomposing soil organic matter. Before sowing, inorganic fertilized soil had the highest soil respiration value, likely due to the presence of easy-available nutrients, which could have initially enhanced the activity of the microorganisms. In fact, soil respiration was quickly reduced after three months of sowing. In the presence of plants, however, organic amendments resulted to be more effective in the enhancement of soil respiration. It is likely that *Melilotus officinalis* exudates from roots together to a higher content of labile organic matter in the soil after compost addition enhanced microbial activity and organic matter decomposition. Sun and co-workers (2018) reported that greater root biomass, also related to higher soil water content, contributed to higher soil respiration values in a grassland ecosystem. Also in our case, treatments in which plants had higher root biomass presented higher soil respiration values.

Soil C contents showed no response to the different soil treatments whereas soil N responded to all fertilization treatments but to a different extent, regardless the presence of plant. The organic amendment did not increase soil C content probably because the microorganisms used the most available Carbon fraction as their activity was particularly stimulated by the rhizosphere, as demonstrated by soil respiration values. The enhancement of microbial activity with the input of organic matter has been shown in several studies (Guerrero et al., 2007; Bustamante et al., 2010).

The significant increase of soil N in organic amendments compared with Inorganic fertilizer may be due to leaching of N in inorganic fertilizer-treated pots (Jing et al., 2017). Also, Adekiya et al., 2020, observed that soil N was higher in organic-amended soils with poultry manure than in an NPK fertilized soil. Organic amendments release N gradually, whereas N from inorganic fertilizers can be more easily lost by volatilization, leaching and/or denitrification. On the other hand, Escanhoela and co-workers (2019), hypothesized that the increases in soil N stocks after poultry litter addition could be related to degraded organic matter allocated in the soil profile. They also showed that organic farming increased N but not C soil stocks compared with conventional systems. The same behaviour was observed for soil K and Na before sowing, they were lower in inorganic fertilized soils.

### Responses of plants to soil amendment treatments

Germination patterns are modified by external (environmental) and internal factors (Rout et al., 2000). In our study, seed germination was clearly affected by soil treatments. The highest germination percentage was observed in high compost-amended soils (**Figure 2**), followed by inorganic fertilized soils. These soils presented also the highest emergency and germination rates. These parameters, however, were very similar in control than in C-Low treatment.

Enhancement of seed germination in the C-high soil might be attributed to the higher availability of macronutrient and micronutrients for the metabolic activity of the germinating seeds (Martinez-Ballesta et al., 2020). The application of a high dose of compost probably has also improved the aeration, porosity and water-holding capacity of the soil, characteristics that also boost seed germination (Masaka and Khumbula, 2007). Usmani and co-workers (2018) also reported that increased applications of vermicomposted fly ash enhanced seed germination of *Lycopersicon esculentum* and *Solanum melongena*.

Concerning the increased and quicker seed germination in inorganic fertilized soil, it may be ascribed to the presence of easy-available nutrients, including nitrogen in the form of nitrate. The fertilizer used in this study is an ammonium nitrate-based fertilizer. Nitrate is known to function not only as a nutrient but also as a signalling molecule acting in seed germination (Duermeyer et al., 2018). Indeed, it has been shown that nitrates stimulate seed germination at low concentrations in several plant species such as *Arabidopsis* (Alboresi et al., 2005) and barley *Hordeum distichum* (Wang et al., 1998). In this regard, also Mohammadi and co-workers (2013) observed higher germination percentages with increasing fertilizer concentrations.

From this study, it was clear that soil amendments, especially compost, have enhanced plant growth. The highest biomass values of *M. officinalis* grown in organic-amended soil at both doses may be a result of the highest release of essential nutrients from the organic fertilizer. In fact, whereas inorganic fertilizers are immediately available for plants, releasing nutrients during a short period, organic amendments are mineralized over time and consequently supply plants with nutrients during a longer period supporting further growth. Lima and co-workers (2004) also observed an increase in the corn plant (*Zea mays* L.) growth after amendment with urban waste compost. On the other hand, Suge and co-workers (2011) showed that organic fertilizer provided a higher yield of *Solanum melongena* than a urea fertilizer.

Leaf N was higher in plants grown in compost-amended soils reflecting soil N contents (that were higher in C-High and C-Low soils). N-fixing plants are less dependent on soil N for their functioning and growth because two different processes supply N: symbiotic fixation and uptake from the soil, in percentages that depend on nodule activity and soil N availability (Peoples and Craswell, 1992). However, N fixation is quite costly energetically and requires a large number of photosynthates (Schulze 2004) as compared with absorbing N from soil (Kaschuk et al., 2009). In this context, our results have shown that with the same assimilation rates, compost-amended plants have grown more than control and inorganic fertilized ones. This fact may be an indication that Control and Inorganic fertilized plants invested more assimilates in N fixation and uptake than the organic-amended plants that had more available N in the soil at the end of the experiment. Indeed, the increase of available N in soil reduces N_2_ fixation rates (Cabeza et al., 2014). Also, P was higher in compost-amended plants than in the other cases. There is a tight relation between N and P uptake in plants. Schleuss and co-workers (2020) found that in grassland ecosystems the combined addition of N and P enhanced the accumulation of soil organic P, the arbuscular mycorrhizal fungi gene abundance and P plant uptake compared to the addition of only P. On the other hand, it has been reported that in low-P soils, the legume white lupin showed an increased N content in shoots and roots as a response to P fertilization (Schulze et al., 2006).

Together with an increase in leaf N and P, leaves from compost-amended plants presented lower levels of K, Ca, Mg and Mn than control and inorganic fertilized plants. This can be due to the formation of complex compounds that make these macroelements unavailable to plants. During the aerobic composting of organic waste, organic matter degrades, and aggregates to produce more stable materials: humic substances (Jurado et al., 2015; Guo et al., 2019) that are incorporated into the soil with compost (Nguyen and Shindo, 2011) forming humus. Humus is rich in functional groups like carboxyl and hydroxyl groups (Piccolo et al., 2019) that can form complexes with cations (Gusiatin and Kulikowska, 2015).

As explained before, we could not observe any difference in CO_2_ assimilation rates and/or stomatal conductance among plants grown on the different soils, despite the diverse leaf N and P levels. Photosynthetic capacity has been reported to be strongly linked to leaf nitrogen because plants invest a considerable amount of leaf nitrogen in Rubisco (Luo et al., 2021). However, this correlation fails in Leguminosae plants as they can fix atmospheric N_2_. In our study, any plant suffered N deprivation. Indeed, it has been shown that nitrogen concentration per unit leaf mass for nitrogen-fixing plants in nonagricultural ecosystems is universally greater (43–100%) than that for other plants (Adams et al., 2016). Also, leaf P has been shown to influence the photosynthetic capacity of plants but as with N, this happens when P content in soils is limited. Several studies reported lower photosynthesis rates values for plants growing on low-P soils (Reich et al., 1994; Denton et al., 2007) or photosynthetic enhancement by P fertilization on P-limited soils (Cordell et al., 2001). Low leaf P content can decrease the photosynthetic capacity by hindering ATP and NADPH synthesis and ribulose-1,5-bisphosphate regeneration (Campbell and Sage 2006; Chu et al., 2018).

The untargeted metabolomics analysis using FT-ICR MS allowed us to characterize the leaves’ metabolome of plants grown under different fertilization treatments and to differentiate them. Compounds belong to different superclasses, with the lipids and lipid-like molecules being the most represented one. Within the lipids’ classes, fatty acyls are clearly the most abundant molecules in these samples. Bioactive compounds already described in *Melilotus* plants were found in all samples, including coumarin, melilotoside, o-coumaric acid or p-coumaric acid, as well as carvone. However, differences in the metabolome were found in plants grown under the different soil treatments. Gargallo-Garriga and co-workers (2017) have previously reported significant metabolome differences between rainforest tree species in response to changes in nutrient availability, specifically N, P and K. In particular, *Tetragastris panamensis and Alseis blackiana showed* larger investment in protection mechanisms with increasing P availability whereas *Heisteria concinna* increased its primary metabolism with increasing N availability. The 5 most important compounds for separation among the *Melilotus* plants were oleamide, palmitic acid (hexadecanoic acid), stearic acid (octadecanoic acid), 3-hydroxy-cis-5-octenoylcarnitine, and 6-hydroxynon-7-enoylcarnitine. All these were found in lower levels in the C-Low plants. Oleamide, palmitoleamide and linoleamide, amide derivatives of the corresponding fatty acids, were found in higher levels in C-High plants, which presented a higher root and aerial biomass. The functional role of these amides in plants is yet unknown, but a possible function has been attributed to plant growth and development regulation (Kim et al., 2013). Palmitic acid and stearic acid are saturated fatty acids that lowered their levels in treated-soils’ grown plants. A decrease in these two fatty acids was previously reported in *Nigella sativa* L. when grown in the presence of bio-chemical fertilizers as well as in *Sesame indicum* with different fertilizer treatments (Moradzadeh et al., 2021). 3-Hydroxy-cis-5-tetradecenoylcarnitine and 6-hydroxynon-7-enoylcarnitine are acylcarnitines that decreased their levels in *Melilotus* plants grown in fertilized soils. Acylcarnitines also decreased their levels in *Pistia stratiotes* plants when the soil was supplemented with copper and zinc nanoparticles (Olkhovych et al., 2016). In plants, acylcarnitines participate in several developmental processes associated to lipid metabolism, particularly related to fatty acid metabolism (Nguyen et al., 2016).

## Conclusions

The use of the organic amendments at high doses implied an improvement of *M. officinalis* germination and growth, not showing significant differences for gas-exchange response at the leaf level. The combination of organic treatment and plant did not produce a clear effect on the soil physicochemical characteristics during the short experimental period, but a significant enhancement of the soil microbial activity was found.

On the other hand, the metabolite profile of *Melilotus* plants showed differences in plants grown under the different soil treatments, although the compounds already described for this plant as bioactive compounds (coumarin, melilotoside, o-coumaric acid or p-coumaric acid and carvone) were found in all samples. Interestingly, the control group had the highest number of exclusive metabolites, and both PCA and hierarchical clustering showed a clear separation between the control samples and the soil-treated samples. Moreover, some interesting metabolites such as saturated fatty acids, palmitic acid and stearic acid, and acylcarnitines, hydroxy-cis-5-tetradecenoylcarnitine and 6-hydroxynon-7-enoylcarnitine, lowered their levels in plants grown on fertilized-soils. In general, the 5 most important compounds for separation among the *Melilotus* plants were oleamide, palmitic acid, stearic acid, 3-hydroxy-cis-5-octenoylcarnitine, and 6-hydroxynon-7-enoylcarnitine.

Thus, this study provides new insight into metabolic changes in melilotus plants responding to different fertilizing treatments. These changes may have implications in the cultivation of Melilotus plants for medicinal purposes in soils where fertilization is mandatory. In poor soils, fertilization with organic amendment together with planting of melilotus may help to the recovery of soil fertility.

## Material and Methods

### Characteristics of the soil and organic amendments

The soil used in this study was obtained from the surface layer (0–20 cm) of a semiarid agricultural area, abandoned for a long time and occasionally used for grazing, in Montelibretti, a town in the Metropolitan City of Rome, in Lazio (Italy). The soil was air-dried before being passed through a 2 mm sieve. The soil from this site is a sandy textured soil, that is slightly alkaline (pH = 7.6), and has low salinity (0.10 dS/m) and very low contents in total organic carbon (0.75% C) and organic matter (0.74%), these contents being lower than 1%, so it can be considered as a semiarid soil. The compost used in this study was produced at the Maccarese composting plant (AMA Srl.), which manages the organic fraction from the selective collection of the municipal solid waste generated in the municipality of Rome (Italy). This compost showed a pH of 7.9, and an EC of 3.6 mScm^-1^, contents of 44.6% of total organic carbon, 2.67% of total nitrogen and a TOC/TN ratio of 16.7, below the value established for compost maturity (TOC/TN < 20) (Bernal et., 2009). The compost from AMA meets the requirements set by the Italian national Law on Fertilizers for agricultural use and has been certified with the quality trademark of the CIC (Consorzio Italiano Compostatori).

### Experimental design

Polyethylene pots with a diameter of 12 cm were filled with soil mixed thoroughly with compost at two doses (expressed on a fresh mass basis): a) Low dose (Low) comprised 3.85 g of compost per kg of soil, corresponding to a dose of 10 t/ha and b) high dose (High) comprised 7.69 g of compost per kg of soil, equivalent to a dose of 20 t/ha. Positive and negative control treatments were included in the experimental design: the positive treatment (Inorganic) was obtained with the application of an inorganic fertilizer composed by NPK in the proportion of 100:60:73. This was obtained by adding 96 mg kg^-1^ soil of the commercial fertiliser Nitrophoska top 20 (NPK = 20:5:10), and 26 mg kg^-1^ of monopotassium phosphate (NPK = 0:52:34). The negative control treatment (Control) consisted of soil without the addition of fertiliser. The NPK ratios of the amended soils were comparable with soils fertilised with inorganic fertiliser (NPK of 100:60:73). Each treatment was replicated ten times. Before sowing, pots were maintained for six months under greenhouse conditions and under a regular watering regime.

Six months after soil preparation, ten *Melilotus officinalis* seeds, previously subjected to chemical scarification for ten minutes with sulfuric acid 98%, were sowed in each pot for germination trials. Germination of seeds was followed for 30 days, then, only two seedlings per pot were maintained and the remaining were thinned out. The remaining seedlings were of similar length. All pots (n = 40) were distributed in a randomised complete block design and maintained for three months in a greenhouse. Watering of the pots was performed regularly and maintained at 50% of field capacity throughout the experiment. After three months, the plants were harvested, aerial part and roots where weighted, measured and the plant foliar tissues were collected (twenty replicates per treatment), weighed to determine fresh biomass and washed with deionised water to remove any attached particles. Then, one plant per pot was dried in an air-forced oven at 60°C reaching the weight stability The dried samples were weighed again for the determination of water content and dry mass, and then ground to a mean size of 0.5 mm for later analyses. The second plant of each pot was immediately frozen in liquid nitrogen and stored at -80 °C until further analysis. Soil samples were also collected (ten replicates per treatment) and divided into two aliquots. One aliquot was used for the determination of soil respiration. The other one was dried in an air-forced oven at 60 °C reaching the weight stability for the rest of the analyses. Soil samples were collected at the beginning (0 days, just before seed sowing) and at the end of the greenhouse experiment (90 days). Plant samples were collected at the end of the greenhouse experiment (90 days).

### Analytical determinations

The pH and electrical conductivity (EC) of the soil samples were measured in extracts with a 1:2.5 and 1:5 soil:water (*w*/*v*) ratio, respectively (Bustamante et al., 2007). Soil respiration was determined according to the methods of Stotzky (1965) and Anderson (1982).

The following determinations were performed on the soil and leaf samples: 1. Soil and leaf total C and total N were determined in an automatic elemental microanalyzer (EuroVector Elemental Analyser, Milan, Italy); 2. Total P, K, Ca, Mg, Na and heavy metals (Fe, Cd, Cu, Mo, Ni, Pb, Zn, Mn) in leaves and soil were determined in the HNO_3_–HClO_4_ digestion extract by inductively coupled plasma atomic emission spectroscopy (ICP-AES, Shimadzu 9000). All the analyses were conducted in five samples per treatment (n=5).

### Gas-exchange measurements

Net photosynthetic rates (A) and stomatal conductance (g_s_) were recorded using an infrared gas analyser (LI-COR LI-6400) (LI-COR, Lincoln, NE, USA) by enclosing two/three leaves into a 6 cm^2^ cuvette equipped with a 6400-02(B) LED Source. Only mature leaves positioned on the fifth and/or the sixth whorl were selected for measurements. Parameters such as relative humidity, air temperature and photosynthetically active radiation (PAR) in the leaf chamber were recorded simultaneously with gas-exchange data. Measurements were carried out under a 500 µmol s^-1^ molar flow rate, 400 ppm CO_2_, Photosynthetic Photon Flux Density (PPFD) of 1000 mmol m^-2^ s^-1^ and leaf temperature of 30 °C. The relative humidity inside the chamber ranged between 50% and 60%. CO_2_ assimilation (A) and stomatal conductance (g_s_) were calculated by the OPEN 6.3.4 software embedded with the analyzer. Measurements were taken in leaves from five different plants per treatment (n=5) between 10.00 and 16.00 h (solar time). Leaf area was calculated after scanning, using ImageJ software version 1.53t (http://rsb.info.nih.gov/ij/).

### FT-ICR MS

Frozen leaf samples (eight biological replicates per treatment) were lyophilized and pulverized. Metabolite extraction was performed as previously described (Maia et al., 2020). Briefly, plant material (c.a 50 mg) was extracted with 1 mL of 50% methanol/water (LC–MS grade, Merck). Then, the methanolic sample extract was diluted 1000-fold in methanol and analysed by direct infusion on a 7-Tesla SolariX XR Fourier Transform Ion Cyclotron Resonance Mass Spectrometer, equipped with a ParaCell (FT-ICR-MS, Brüker Daltonics). Leucine enkephalin (YGGFL, Sigma Aldrich) was added as internal standard ([M+H]^+^ =556.276575 *m/z*), at a concentration of 0.1 µg/mL, and formic acid (Sigma Aldrich, MS grade) was added at a final concentration of 0.1% (v/v) to all replicates. Spectra were acquired in positive electrospray ionization (ESI^+^), in magnitude mode, with an acquisition size of 4M, and recorded between 100 and 1500 *m/z*, as previously described (Maia et al., 2020). The accumulation time was 0.1 seconds, and 100 transients were accumulated for each spectrum, zero-filled and apodised (half-sine). Online calibration was performed using the monoisotopic *m/z* value of leucine enkephalin. The software Bruker Compass data Analysis 5.0 (Brüker Daltonics, Bremen, Germany) was used to process and retrieve the mass lists, considering peaks with a minimum signal-to-noise ratio of 4. Spectral alignment of the different samples was performed with Metaboscape 5.0 (Brüker Daltonics, Bremen, Germany) using T-ReX 2D (MRMS single spectrum) algorithm. The intensities were normalized by the total spectrum signal. Variables appearing in only one of the fifteen samples and data for the internal standard (leucine enkephalin, neutral mass 555.2693063368 Da) were removed from the dataset.

The number of common and exclusive metabolites for each soil type was calculated and plotted as an intersection plot (Lex et al., 2014). Molecular formulas for each isotopic cluster were assigned using the SmartFormula function in Metaboscape, following a series of heuristic rules proposed (Kind and Fiehn, 2007) and others implemented in SmartFormula (like removing chloride and fluoride containing formulas). Metabolite annotation was performed in Metaboscape using the Human Metabolome Database (HMDB, from 27 May 2022; Wishart et al., 2022), PlantCyc (from 9 May 2022; Hawkins et al., 2021), and LOTUS (from 16 September 2022; Rutz et al., 2022), uploaded to MetaboScape 5.0 and considering a maximum deviation of 1 ppm. Compounds were classified according to ChemOnt chemical taxonomy (Djoumbou et al., 2016) and reporting the “superclass”. For lipids, the “class” was also reported. A metabolic map focused on the citrate cycle and the fatty acid biosynthesis and degradation pathways was built with the identified compounds and their relative quantification in the four different sample groups.

### Statistical Analysis

Analysis of variance (ANOVA) was performed with either soil pH, soil EC, soil respiration, soil/leaf C content or soil/leaf nutrient contents as the dependent variable and organic fertilization as the independent factor. The Fisher post-hoc test was used to investigate the significance of different groups of means, considering a probability level of *P* < 0.05.

For the metabolomics data analysis, multivariate statistical analysis was performed in Python, using the modules *pandas* (Reback et. al., 2021), Scikit-learn (Li & Phung, 2011), SciPy (Virtanen et al., 2020) and NumPy (Harris et al., 2020). Plots were produced with matplotlib (Hunter, 2007) and seaborn (Waskom et al., 2020). Missing values were assigned with one-fifth of the minimum intensity for each m/z across all samples and data was Pareto scaled. Two unsupervised methods were applied: Principal Component Analysis (PCA) and sample Hierarchical Clustering (agglomerative, HCA). HCA was performed using the metric Euclidian distance and the Ward variance minimization algorithm as the linkage method. A classifier for discrimination of the plants grown in the four soil types was obtained by building Partial Least Squares Discriminant Analysis (PLS-DA) models, using the scikit-learn library (Li & Phung, 2011). Model accuracy, R2 and Q2 metrics were also estimated by 8-fold stratified cross-validation. The number of components for the PLS-DA models was 6, chosen to minimize the predictive residual sum of squares (maximize Q2). Variable importance in projection (VIP) scores for each variable were used for feature importance scoring and were calculated as a weighted sum of the squared correlations between the PLS-DA components and the original variable. Analyses were carried out using in-house developed software.

## Supplemental data

The following materials are available in the online version of this article:

**Supplemental Table S1**: Detected metabolites in *M. officinalis* plants grown on Control, Inorganic, C-Low and C-High soils, analysed by FT-ICR-MS in positive (ESI+) ion analysis mode (aligned table). Experimental *m/z* is the *m/z* value detected by FT-ICR-MS, considering H^+^, Na^+^ and K^+^ as possible adducts; the neutral mass (Da) is the corrected mass of the compound without adduct; Compound Name (First annotation) is the first name that appears in the database for that *m/z* value; Compound Formula (Smart Formula) is the formula of the compound calculated by Bruker’s Metaboscape Smartformula tool; for each database, HMDB (Human Metabolome Database), PlantCyc, and LOTUS, is shown the compound ID in the database, the name, formula and the number of possible identifications (match count) in that database. The intensities of the compounds in each replica of the four groups is also shown.

## Acknowledgements and Funding

This work has been conducted in the framework of the TA project MOPS financed by the Integrating Activity, and the Research Infrastructure project European Network of Fourier-Transform Ion-Cyclotron-Resonance Mass Spectrometry Centers (EU FT-ICR-MS) funded by the European Union’s Horizon 2020 research and innovation programme, Grant Agreement No 731077. The authors acknowledge the PhD grant 2021.06370.BD to F. Traquete from Fundação para a Ciência e a Tecnologia (Portugal). This work was also supported by the Short-term mobility Program for Scientists/Researchers from Italian and Foreign Institutions (2020) of the National Research Council of Italy (CNR) of Dr. Nogués (prot. CNR 0060457/2020 dated 01/10/2020).

## Data availability

The metabolomics data that support the findings of this study are available in figshare data repository with the identifier https://doi.org/80.10.6084/m9.figshare.22557493 (Nogues et al., 2023).

